# Non-autonomous induction of epithelial lineage infidelity and hyperplasia by DNA damage

**DOI:** 10.1101/630954

**Authors:** Lindsey Seldin, Ian G. Macara

## Abstract

Several epithelial tissues contain stem cell reserves to replenish cells lost during normal homeostasis or upon injury. However, how epithelial tissues respond to distinct types of damage, and how stem cell plasticity and proliferation are regulated in these contexts, remain poorly understood. Here, we reveal that genotoxic agents, but not mechanical damage, induce hyperplasia and lineage infidelity in three related epithelial tissues: the mammary gland, interfollicular epidermis and hair follicle. Furthermore, DNA damage also promotes stromal proliferation. In the mammary gland, we find that DNA damage activates multipotency within the myoepithelial population and hyper-proliferation of their luminal progeny, resulting in tissue disorganization. Additionally, in epidermal and hair follicle epithelia, DNA damage induces basal cell hyperplasia with the formation of abnormal, multi-layered K14+/K10+ cells. This behavior does not involve apoptosis or immunity, and is epithelial cell non-autonomous; stromal fibroblasts are both necessary and sufficient to induce the epithelial response. Thus, genotoxic agents that are used chemotherapeutically to promote cancer cell death can have the opposite effect on wild-type epithelial tissue, paradoxically promoting hyperplasia and inducing both stemness and lineage infidelity.

## INTRODUCTION

The mammary epithelium arises from the embryonic epidermis, and is composed primarily of two distinct layers, an outer (basal) layer of contractile myoepithelial cells positive for keratin 14 (K14+) and an inner layer of luminal cells positive for keratin 8 (K8+). Lineage-tracing has shown that although the embryonic mammary gland develops from bipotent stem cells, postnatally each cell type is maintained by distinct unipotent progenitors^1–4^. Strikingly, however, the myoepithelial cells become highly plastic when dissociated from their native tissue^1, 2, 5–9^, such that a single transplanted cell can regenerate a complete mammary ductal tree within the cleared fat pad of a recipient mouse^6, 7^. A similar loss of fate restriction has been observed in the skin. Stem cells within the bulge of the hair follicle do not normally give rise to the infundibulum, interfollicular epidermis or sebaceous gland, but lose this restriction when isolated and engrafted onto a recipient mouse, where they can regenerate the entire skin epithelium^10^. Since this potential for multipotency is efficiently suppressed in unperturbed tissue, it begs the question of why such a property has been retained.

One possibility is that a reversion to stemness is needed in response to tissue damage, as a mechanism of wound healing. There is precedent for such a function in the skin, intestine and lung. Hair follicle stem cells, for example, are rapidly recruited to the interfollicular epithelium after wounding, and function in short-term repair, though few contribute to the tissue over the longer-term^11, 12^. Moreover, if bulge cells are destroyed, hair germ cells can colonize the empty niche and mediate hair regeneration^13^. Similarly, in intestinal crypts, ablation of Lgr5+ stem cells triggers quiescent Bmi1+ cells to convert into Lgr5+ cells to maintain gut homeostasis^14^; and in the lung, ablation of tracheal basal stem cells can trigger committed secretory cells to de-differentiate and replace them^15^. Therefore, we considered that the multipotency potential of mammary myoepithelial cells, identified through their isolation and transplantation, might represent a damage response.

## RESULTS

### DNA damage promotes multipotency in mouse mammary myoepithelial cells in situ

We first confirmed that K14+ myoepithelial cells are unipotent in their unperturbed in situ environment during morphogenesis, as reported previously^1–4^. We used a doxycycline-inducible mouse model to perform saturation lineage-tracing of myoepithelial cell progeny (K14-rtTA; TetO-Cre; tdTomato^fl/wt^) (Fig.1A)^16–18^. To promote tdTomato expression, we maintained the mice on doxycycline chow for six weeks from puberty onset (three-weeks old) before sacrifice and analysis (Fig.1B). We performed fluorescence-activated cell sorting (FACS) analysis to determine the efficiency of tdTomato-positive myoepithelial cell labeling, as well as potential leakage into the luminal population. We found an average of 81.8% tdTomato-positive myoepithelial cells and 0.9% tdTomato-positive luminal cells following the six-week treatment period, indicating that this transgenic system has very low leakage rates and successfully labels the majority of myoepithelial cells over this timeframe (Fig.S1V). We also used a tamoxifen-inducible transgenic mouse model to lineage-trace K8+ luminal mammary cell progeny by injecting tamoxifen at puberty onset (K8-CreER; tdTomato^fl/wt^) (Fig.S1S-T)^1^. Both myoepithelial and luminal cells were unipotent, corroborating published data (Figs.1D, S1U).

**Figure 1.**
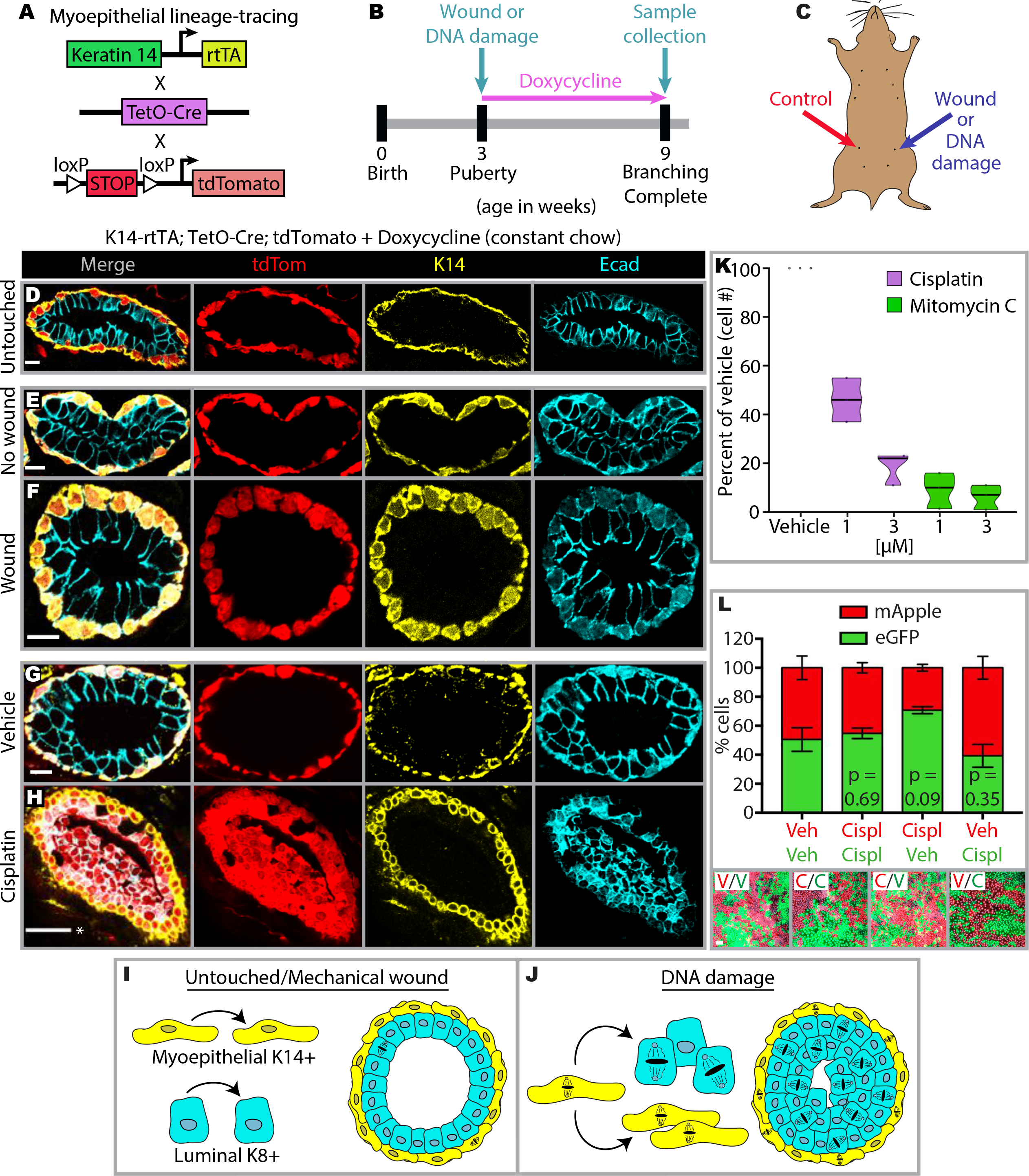
DNA damage promotes multipotency in mouse mammary myoepithelial cells in situ. A) Diagram illustrating the three transgenic mouse lines crossed for mammary myoepithelial lineage-tracing. B) Experimental timeline of doxycycline-driven induction of myoepithelial cell labeling. C) Treatments were injected into mammary fat pads at puberty onset, as illustrated. D) Immunofluorescence images revealing that unperturbed myoepithelial cells are unipotent, giving rise to myoepithelial progeny throughout morphogenesis (3/3 mice, 100%). E-F) Immunofluorescence images showing that myoepithelial cells maintain unipotency throughout morphogenesis in both unwounded (E) and mechanically-wounded (F) mammary glands (3/3 mice per condition, 100%). G-H) Immunofluorescence images illustrating that, unlike vehicle (G), the DNA interstrand crosslinking agent Cisplatin (H) promotes myoepithelial cell multipotency, which results in luminal filling (3/3 mice per condition, 100%). I-J) Cartoon illustration of mammary cell plasticity in unperturbed and mechanically-wounded mammary glands (I) versus DNA-damaged mammary glands (J). K) Violin plots reveal the effect of the DNA crosslinking agents Cisplatin and Mitomycin C on cell death in cultured EpH4 mouse mammary epithelial cells (Cisplatin vehicle - PBS; Mitomycin C vehicle - dH_2_O) (n=3 experimental replicates). L) Bar graphs of results from cell mixing experiments using stably transduced mApple and eGFP EpH4 sorted cell lines, showing that surviving Cisplatin-treated (“Cispl”, “C”) cells do not have a proliferative advantage over vehicle (PBS)-treated (“Veh”, “V”) cells when plated at a 1:1 ratio, and vice versa (n=3 experimental replicates). Scale bars are 50 *μ*m (*) or 10 *μ*m.

To test our hypothesis that tissue damage promotes mammary cell plasticity, we introduced two distinct types of damage in vivo: mechanical wounding or genotoxicity. We used one mammary fat pad for the experimental condition, and another in the same mouse as an internal control (Fig.1C). For mechanical wounding, we made deep lacerations throughout the mammary tissue using a scalpel. Alternatively, we injected a DNA crosslinking agent directly into the mammary fat pad. We performed these interventions on three-week old pubertal mice, then followed the same feeding and analysis regimen described above (Fig.1B). Surprisingly, mechanical wounding had no impact on mammary architecture or myoepithelial cell behavior, as determined by lineage-tracing, and their unipotency was maintained (Figs.1E-F, S2A-D). However, unlike vehicle, DNA damage (induced by the chemotherapeutic agents Cisplatin or Mitomycin C) triggered a remarkable disruption of myoepithelial lineage specification (Figs.1G-H, S1A-R). Large numbers of tdTomato+ cells derived from the myoepithelium were generated (Figs.1H, S1B-J, L-R), positive for the luminal marker E-cadherin (Fig.1H). Furthermore, this unexpected response to DNA damage resulted in the perturbation of normal epithelial architecture, as the tdTomato+ cells had displaced into and filled up the lumens (Figs.1G-H, S1A-R, S2A-B, E-H). Notably, this luminal filling resembles ductal carcinoma in situ, an early pre-invasive stage of breast cancer^19, 20^.

In contrast with these in vivo findings, treating EpH4 mammary epithelial cells in culture with Cisplatin or Mitomycin C caused cell death, as expected (Fig.1K)^21^. To determine whether the in vivo-specific response could be a consequence of Cisplatin resistance that endows cells with a proliferative advantage, we tested if EpH4 mammary cells that survived Cisplatin treatment could outcompete vehicle-treated cells in culture. We established EpH4 cell lines expressing mApple or eGFP, and separately incubated each with either Cisplatin or vehicle. Surviving cells from each treatment were then mixed at a 1 (Cisplatin-resistant): 1 (vehicle) ratio, and color ratios were analyzed 48 hr later. We found no significant differences between the percentages of Cisplatin-resistant vs. vehicle-treated cells, suggesting that Cisplatin resistance does not underlie the in vivo mammary response to DNA damage (Fig.1L).

Taken together, these experiments reveal that DNA damaging agents, unlike mechanical wounding, cause a paradoxical hyperplasia and lineage infidelity in wild-type mammary epithelium in situ (Fig.1I-J). The striking differences between mammary cell responses in vivo and ex vivo suggest that the tissue microenvironment may play a critical role in the tissue response to DNA damage.

### DNA damage promotes hyper-proliferation and lineage infidelity in intact mouse epidermis

We next asked if these responses to DNA damage are mammary-specific or generalizable to other epithelial tissues. The mammary gland is a sub-appendage of the epidermis, as mammary placodes first emerge from epidermal epithelium during embryogenesis. However, the two tissues diverge in architecture and cell lineage pathways. K14+ basal cells of the epidermis are bipotent, balancing between self-renewal and production of non-proliferative K10+ suprabasal cells that terminally differentiate to form an impenetrable skin barrier^22^. Notably, basal cells in adult epidermis have very low proliferation rates (Fig. 2D, BrdU channel). To determine how the epidermis responds to mechanical damage, we scraped the backskin of 3-6 week old wild-type adult mice, and analyzed the wounded skin region after one week. We found no significant alterations in epidermal cell organization, tissue thickness or proliferation (based on BrdU+ staining), compared to untouched, age-matched control tissue (Fig.2A, D-E). However, one week after Cisplatin ‘painting’ onto shaved and scraped backskin (Fig.2B), the gross tissue exhibited a rough, scabbed and scaly appearance compared to vehicle (Fig.S3Q-R). To test for DNA damage, we stained skin sections for γH2AX and detected positive cell nuclei within the basal layer 1 day post-treatment, as compared to control tissue, which was γH2AX-negative (Fig.S3G-H). Notably, a significant number of dermal cells also stained positive for γH2AX following Cisplatin treatment (Fig.S3G-H).

**Figure 2.**
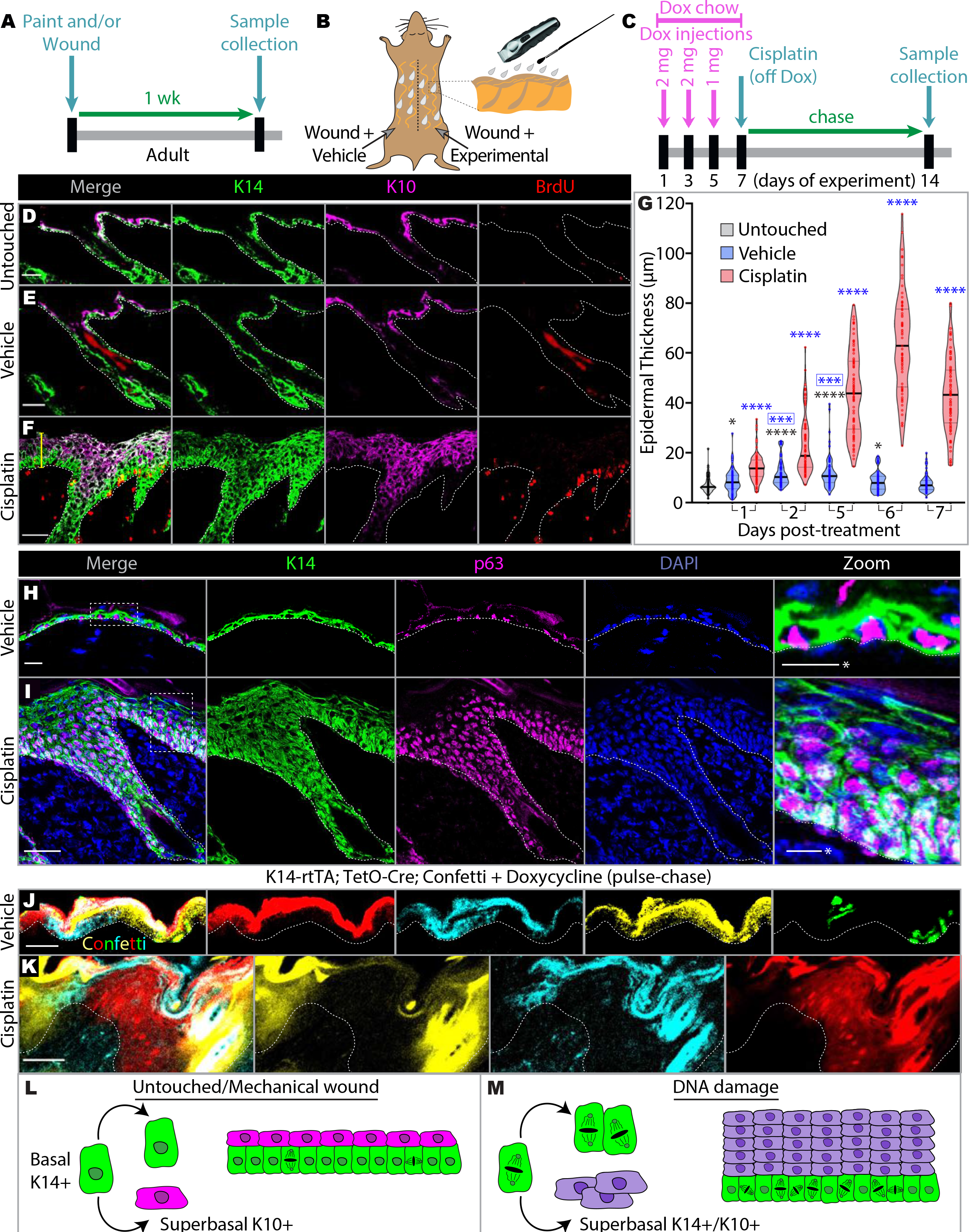
DNA damage promotes hyper-proliferation and lineage infidelity in intact mouse epidermis. A) Experimental timeline of drug treatment on wild-type C3H mouse shaved backskin. B) Cartoon illustrating the wounding (“scraping”) of mouse backskin by creating abrasions with a razor, followed by vehicle or drug painting. The shaved backskin regions for vehicle and experimental drug application are separated by a tuft of unshaven hair (dotted line, see photograph in Fig.S3P). C) Experimental pulse-chase timeline of doxycycline-driven induction of basal cell labeling in K14-rtTA; TetO-Cre; Confetti mice. D-E) Immunofluorescence images showing that mechanical wounding of backskin (E) does not cause large changes compared with untouched control tissue (D). F) Immunofluorescence images revealing that Cisplatin treatment in combination with wounding promotes hyper-proliferation in the basal and dermal cell compartments, as well as epidermal lineage infidelity and tissue thickening. The yellow bracket represents the region measured to determine “epidermal thickness”, as quantified in (G). G) Violin plots of epidermal thickness over 7 days following vehicle and Cisplatin treatment (n=3 mice per violin). Black * is vehicle compared to untreated; blue * is Cisplatin compared to same day vehicle; boxed blue * is vehicle compared to day 1 vehicle. *p < 0.05, **p < 0.001, ***p < 0.0001, ****p<0.00001. H-I) Immunofluorescence images of p63 antibody staining in vehicle- (H) and Cisplatin- (I) treated epidermis. J-K) Immunofluorescence images of lineage-traced K14-rtTA; TetO-Cre; Confetti mouse backskin, treated according to the experimental timeline in (C), reveals clonal dynamics upon vehicle (J) and Cisplatin (K) treatment (see additional images in Fig.S3I-O) (n=4). L-M) Cartoon illustration of cell behavior in unperturbed or mechanically-wounded epidermis (L) versus DNA-damaged epidermis (M). Scale bars are 10 *μ*m (*) or 50 *μ*m.

To probe for a proliferative response to Cisplatin, we used BrdU pulse-labeling. Tissue analysis 7 days post-treatment revealed a substantial increase in BrdU+ basal cells compared with vehicle (PBS)-treated controls (Fig.2E-F). Many dermal cells also showed BrdU labeling following Cisplatin treatment (Fig.2E-F). Expression of the apoptotic marker cleaved caspase-3 was absent from tissue both 1 and 2 days post-treatment, suggesting that these responses were not a compensatory reaction to apoptosis (Fig.S3A-D).

Strikingly, Cisplatin treatment caused a massive expansion of the suprabasal layers, with the majority of cells staining double-positive for K14 and K10, a state that is only rarely detected in normal epidermis (Fig.2E-F). The stem cell marker p63, normally confined to the basal cell layer, was inappropriately expressed throughout these expanded layers despite their lack of BrdU+ labeling, suggesting defective differentiation (Fig.2H-I). Nonetheless, these cells maintained robust basolateral expression of E-cadherin, demonstrating retention of their polarity and epithelial identity (Fig.S3T-U). The Cisplatin-treated tissue also appeared to be under stress, as indicated by positive keratin 6 (K6) staining in the interfollicular epidermis that is entirely absent from control tissue (Fig.S3V-W).

Importantly, Cisplatin-induced hyper-proliferation and lineage infidelity were not dependent on hair follicle stage, as DNA-damaged skin containing hair follicles in either the growth (anagen) or resting (telogen) phases of the hair cycle exhibited similar interfollicular epidermal responses (Figs.2E-F, S3X-Y). These DNA damage effects remained localized to the treatment area, since vehicle and Cisplatin treatments were applied to separately shaved backskin regions of the same mouse, partitioned only by a tuft of hair (Fig.S3P).

The increase in tissue thickness of Cisplatin-treated skin peaked between 5 - 7 days post-treatment (Fig.2G). This response then waned over a period of 5 weeks (Fig.S4A). In addition, backskin exposed to a second round of Cisplatin treatment was also able to resolve back to its original thickness (Fig. S4B). These observations suggest that the epithelial response to Cisplatin is reversible, and that no oncogenic transformation occurs, at least over the short term. We conclude that hyper-proliferative basal cells activated by Cisplatin eventually lose their proliferative capacity, apoptose, and/or undergo differentiation and are ultimately shed.

To confirm that the massively expanded suprabasal population came from the hyper-proliferative basal cells, we performed pulse-chase experiments using the K14-rtTA; TetO-Cre; tdTomato lineage-tracing mouse line described above (Fig.1A). This pulse-chase approach was necessary to ensure that only the progeny of cells originating in the basal population would be traced, since all cells following Cisplatin treatment express K14. We both fed and injected mice with doxycycline over a one-week period, then transferred them to a normal diet after Cisplatin painting, and collected samples one week later (Fig. 2C). While tdTomato expression in vehicle-painted backskin was confined to the basal layer, all cell layers in the Cisplatin-painted backskin retained the tdTomato label, thus confirming that the expanded suprabasal population was generated from hyper-proliferative basal cells as opposed to pre-existing suprabasal cells (Fig. S3E-F). We also performed the same experiment using multicolor Confetti mice (K14-rtTA; TetO-Cre; Confetti), which generate stochastic labeling of myoepithelial cells with one of four distinct colors^23, 24^. These analyses corroborated that the expanded layers arise from hyper-proliferating basal cells, while also revealing the clonal dynamics during this overgrowth period (Figs.2J-K, S3I-O)^25, 26^. Distinct clones originated from both the interfollicular epidermis and hair follicles. Consistent with previous work, Confetti lineage-tracing revealed large segments of alternating colors, suggesting that the interfollicular epidermis is sub-compartmentalized into segregated regions with suprabasal layers generated from individual stem cells interspersed throughout the basal layer^11^. We conclude that Cisplatin treatment either activates a specific subpopulation of basal cells that are primed to hyper-proliferate, or stochastic clonal drift allows some clones to outcompete others. Furthermore, these lineage-tracing findings confirm that hyper-proliferation precedes lineage infidelity, as Cisplatin-treated basal cells retained K14-only expression similar to untreated tissue, while generating all of the suprabasal layers that exhibit lineage infidelity.

In contrast, Cisplatin treatment of cultured primary epidermal cells did not induce either lineage infidelity or hyper-proliferation, showing that the in vivo environment is required for this response (Fig.S3S). Together these data demonstrate that multiple wild-type epithelial tissues with otherwise low proliferation rates undergo hyper-proliferation, lineage infidelity and defective tissue organization upon treatment with genotoxic chemotherapeutic agents (Figs.1I-J, 2L-M).

### The epidermal response to DNA damage is cell non-autonomous

The contrary responses of mammary and epidermal cells to genotoxic agents in vivo versus in culture argues strongly for involvement of the tissue microenvironment in modulating epithelial cell behaviors (Figs.1K, S3S). It is well-established that epithelial tissues are influenced by constituents of their microenvironment, which include fibroblasts, adipocytes, immune cells, and vasculature. Stromal components of the mammary fat pad are essential for promoting mammary gland morphogenesis and remodeling^27, 28^; epidermal wound healing requires dermal fibroblast recruitment into the wound bed; and hair follicle morphogenesis depends on extensive epithelial-mesenchymal crosstalk^29^. Notably, microenvironmental signals can also promote epithelial tumorigenesis. Cancer-associated fibroblasts (CAFs) have been linked to the progression of both triple-negative breast cancer and basal cell carcinoma^30–32^. To test whether DNA damage to stromal cells can elicit the epithelial behaviors observed in vivo, we first injected Cisplatin under the skin epithelium directly into the dermis (“intradermal injection”), and analyzed the tissue one week post-injection (Fig.3C). Similar to our previous scraping and painting approach, this treatment induced both dermal and epithelial hyper-proliferation, causing significant epidermal thickening and lineage infidelity compared with vehicle (Fig.3A-B, D). These data indicate that DNA damage alone, in the absence of wounding, is sufficient to promote a response. Notably, intradermal injections promoted hyper-proliferation and lineage infidelity down the outer root sheath of the hair follicle, with the follicular epithelium expressing both K14 and K10 (typically absent from normal hair follicles) (Fig.3A-B). Furthermore, the bottommost region of the anagen hair follicle (normally composed of proliferative BrdU+ matrix cells that give rise to all inner layers of the differentiated hair shaft) showed no BrdU labeling, and also co-stained positive for both K14 and K10 following Cisplatin but not vehicle intradermal injections (Fig.3G-H). This effect was exclusive to intradermal injections, as the localized painting of Cisplatin onto scraped backskin did not allow the drug to penetrate deep enough to promote this effect at the hair follicle base (Fig.3E-F).

**Figure 3.**
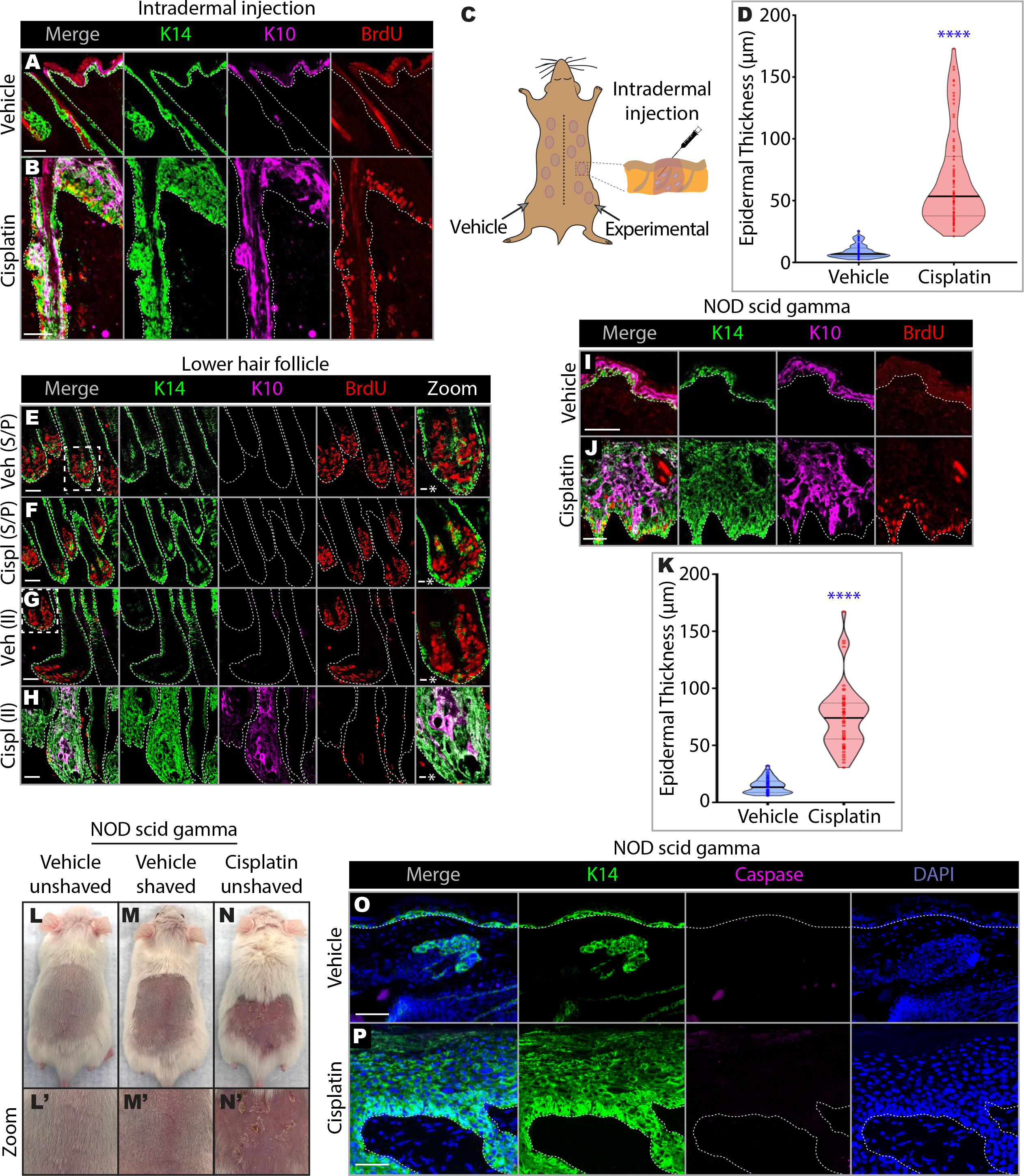
The epidermal response to DNA damage is cell non-autonomous. A) Immunofluorescence images revealing that intradermal injections of Cisplatin (B) but not vehicle (A) promote hyper-proliferation in the interfollicular, follicular and dermal cell compartments, epidermal thickening in the interfollicular epidermis, as well as lineage infidelity in the interfollicular epidermis and hair follicle. C) Cartoon illustrating the injection of vehicle and experimental treatments into the dermal region of shaven mouse backskin. The shaved backskin regions for vehicle and experimental drug injection are separated by a tuft of unshaven hair (dotted line, see photograph in Fig.S3P). D) Violin plots of epidermal thickness 7 days following vehicle and Cisplatin intradermal injections (n=3 mice per violin). ****p<0.00001. E-F) Immunofluorescence images showing that scraping and painting (“S/P”) vehicle (E) or Cisplatin (F) onto shaven backskin does not cause changes in proliferative matrix cells of the lower hair follicle. G-H) Immunofluorescence images showing that, compared with the proliferative matrix cells present following intradermal injection (“II”) of vehicle (G), no proliferative matrix cells are present following intradermal injection of Cisplatin (H). Instead, epithelial cells at the hair follicle base demonstrate lineage infidelity, with double-positive K14/K10 expression. I-J) Immunofluorescence images of epidermis from immunocompromised NOD scid gamma (NSG) mice treated with vehicle (I) or Cisplatin (J). Cisplatin causes hyper-proliferation, epidermal thickening and lineage infidelity, comparable to its effect on wild-type C3H mouse backskin that has intact immunity. K) Violin plots of NSG epidermal thickness 7 days following vehicle and Cisplatin treatment (n=2 mice per violin). ****p<0.00001. L-N) Photographs of gross NSG backskin appearance 7 days following vehicle (L-M) or Cisplatin (N) treatment (L’-N’ are corresponding image zoom-ins). O-P) Immunofluorescence images of cleaved caspase-3 staining of NSG epidermal tissue treated with vehicle (O) or Cisplatin (P) reveals no apoptotic cells, suggesting that macrophages and dendritic cells do not clear away dead cells upon Cisplatin treatment. This further confirms that epidermal cells do not undergo apoptosis in response to DNA damage. Scale bars are 10 *μ*m (*) or 50 *μ*m.

To identify the dermal component(s) promoting these epithelial behaviors, we took a candidate approach. Psoriasis is a chronic, inflammatory skin disease caused by overactive innate and adaptive immune responses, and psoriatic skin, like Cisplatin-treated skin, exhibits keratinocyte hyper-proliferation, hyperkeratinosis and defective cell differentiation^33^. Therefore, we tested whether immune cells are involved in the epidermal Cisplatin response by treating the backskin of immunocompromised NOD scid gamma (NSG) mice, which lack both adaptive and innate immune function. Contrary to expectations, we found that gross tissue scabbing, hyper-proliferation, tissue thickening, and lineage infidelity still occurred in these mice following Cisplatin treatment, demonstrating that immune cells are not required for the Cisplatin effect (Fig.3I-N)^34, 35^. This particularly striking finding challenges current dogma that immune cells, particularly T cells, are critical for mounting efficient skin responses to injury^36^. Furthermore, since macrophages and dendritic cells phagocytose apoptotic cells, we reasoned that the lack of apoptotic cells observed in wild-type Cisplatin-treated backskin could be due to their rapid clearing by these immune scavengers^37^. After staining Cisplatin-treated NSG backskin (that lack macrophage and dendritic activity) with cleaved caspase-3, however, we still did not detect the presence of apoptotic cells (Fig.3O-P).

### Dermal fibroblasts are necessary and sufficient for the epithelial response to DNA damage

An alternative candidate cell type that could drive the epithelial response to DNA damage is the dermal fibroblast, which can stimulate keratinocyte proliferation, and is important for epidermal morphogenesis, maintenance and regeneration^38, 39^. To determine whether fibroblasts are required for the epithelial response to Cisplatin, we ablated them by pre-treating Col1a2-CreER; DTA (Diphtheria toxin A) mice with 4-Hydroxytamoxifen (4OHT), then treated their backskin with Cisplatin (Fig.4A)^40^. In this system, 4OHT promotes Diphtheria toxin A activation in, and subsequent killing of, Col1a2+ fibroblasts (Fig.4B). Col1a2-CreER; DTA mice pretreated with vehicle and then treated with Cisplatin exhibited a hyper-proliferative response similar to that of wild-type Cisplatin-treated mice, whereas fibroblast ablation with 4OHT caused a dramatic decrease in proliferation, epidermal thickening, and lineage infidelity (Fig.4C-E). We conclude that fibroblasts are necessary to drive the epithelial responses to Cisplatin.

**Figure 4.**
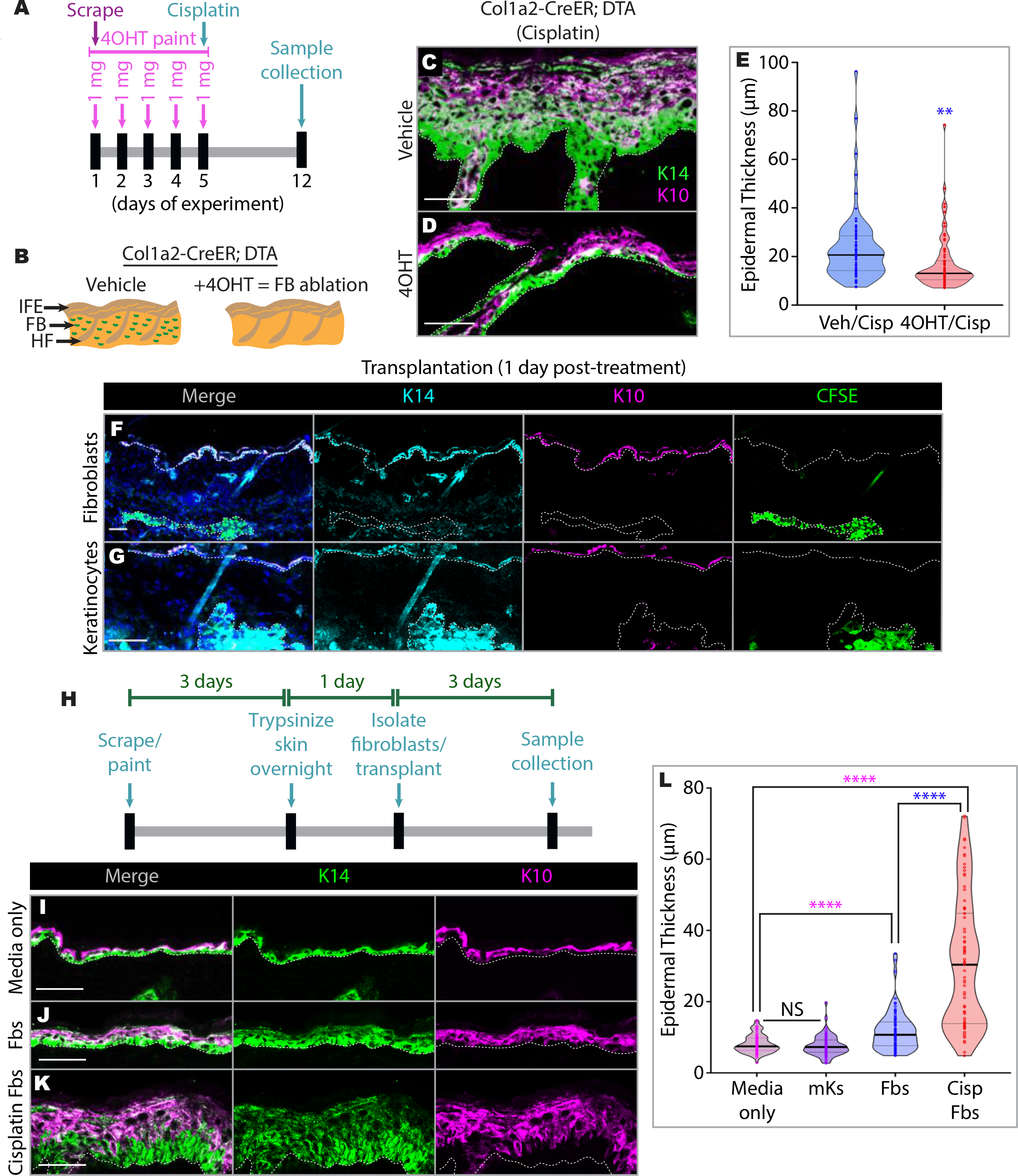
Dermal fibroblasts are necessary and sufficient for the epithelial response to DNA damage. A) Experimental timeline of 4-hydroxytamoxifen (4OHT)-driven induction of dermal fibroblast ablation in Col1a2-CreER; DTA mice prior to Cisplatin treatment. B) Left cartoon illustrates the different compartments of the skin: “IFE” - interfollicular epidermis; “FB” - fibroblast; “HF” - hair follicle. Right cartoon illustrates that painting 4OHT onto Col1a2-CreER; DTA backskin promotes dermal fibroblast ablation. C-D) Immunofluorescence images of Col1a2-CreER backskin, showing that Cisplatin induction of hyper-proliferation and epidermal thickening occurs in vehicle (C) but not 4OHT-treated tissue (D) that lack dermal fibroblasts. E) Violin plots of epidermal thickness in Col1a2-CreER mice following the treatment timeline in Fig.4A (n=3 mice per violin). **p < 0.001. F-G) Immunofluorescence images of CFSE-labeled dermal fibroblasts (F) and keratinocytes (G) 1 day post-transplantation, confirming that both cell types survive and are properly incorporated into the dermis of recipient skin. H) Experimental timeline of transplantations. I-J) Immunofluorescence images of recipient C3H epidermis following media injections (I), or transplantation of untreated (J) and Cisplatin-treated (K) dermal fibroblasts from syngeneic C3H donor mice. L) Violin plots of epidermal thickness in recipient mice following transplantation, according to the experimental timeline in Fig.4H (n=3 mice per violin). Pink * is compared to media only; Blue * is compared to untreated fibroblasts. ****p<0.00001. “mKs” - mouse keratinocytes; “Fbs” - fibroblasts; “NS” - not significant. Scale bars are 50 *μ*m.

Probing further, we performed a converse experiment by transplanting fibroblasts from Cisplatin-treated dermis into the dermis of syngeneic recipients. We first confirmed that transplanted fibroblasts and keratinocytes (used as a negative control) survived post-transplantation and were positioned properly in the recipient dermal region by tracking cells with a CFSE fluorescent label (Fig.4F-G). We then performed separate transplantation experiments using fibroblasts isolated from untreated and Cisplatin-treated mice, and assessed epidermal cell behavior 3 days post-transplantation (Fig.4H). We also transplanted keratinocytes from the same donor mice or injected media as additional controls. Strikingly, the fibroblast transplantations, unlike control keratinocytes and media, were sufficient to trigger hyper-proliferation, tissue thickening and lineage infidelity (Fig.4I-L). Interestingly, the isolation and transplantation of fibroblasts from untreated mice was sufficient to promote some epidermal thickening and lineage infidelity, though to a significantly lesser degree than fibroblasts from Cisplatin-treated mice (Fig.4I-L). This result suggests that the isolation procedure alone can prime the fibroblasts, as has been observed for other cell types^9, 10^. Overall, we conclude that epithelial activation by DNA damaging agents is cell non-autonomous, and that dermal fibroblasts are both necessary and sufficient to drive this response (Fig.S4C-D).

## DISCUSSION

We have discovered a counter-intuitive effect of genotoxic agents on epithelial tissues in vivo. Chemotherapeutic DNA crosslinking agents that block cell proliferation and trigger apoptosis in culture have the opposite effects in the intact skin and mammary gland, promoting epithelial hyperplasia and lineage infidelity. Unexpectedly, this response is cell non-autonomous, but is independent of both innate and adaptive immunity. We found that, instead, dermal fibroblasts are both necessary and sufficient to drive the epithelial response. We propose that in response to DNA damage, the fibroblasts release factors that stimulate epithelial cell hyper-proliferation, which in turn causes defects in lineage specification (Fig.S4C-D). Whether a similar mechanism accounts for lineage switching in the mammary gland remains to be established.

The de-repression of cell plasticity in mammary glands subjected to DNA damage is reminiscent of mammary cell reprogramming into multipotent stem cells upon dissociation from their native tissue^1, 2, 5–8^. Oncogenic activation of unipotent mammary cells in situ can also promote multipotency^41^. Similarly, in both the interfollicular and follicular epidermal compartments, we show that DNA damage promotes hyperplasia and lineage infidelity, underscoring the generalizability of the effect across multiple epithelial tissue types. This response does not require innate or adaptive immunity^42^, but the hyperkeratinosis phenotype is nonetheless similar to that seen in immune and inflammation-triggered skin diseases such as psoriasis and atopic dermatitis, and in experimentally-induced inflammation^43^, demonstrating how multiple cellular mechanisms driven by distinct molecular pathways can converge to cause similar gross tissue pathologies.

The drugs Cisplatin and Mitomycin C used in this study generate interstrand and intrastrand DNA crosslinks that block replication and transcription. They are used therapeutically for a variety of epithelial cancers, by promoting apoptosis of rapidly cycling tumor cells^44–48^. Strikingly, however, aside from work on Cisplatin-induced nephrotoxicity, most studies characterizing the effects of genotoxic agents on cell behavior have been limited to cancer cell lines in culture^21,49^. Although DNA damage was previously reported to cause defective differentiation in cultured mammary epithelial cell lines, the relevance of these ex vivo responses to in vivo mammary tissue is unclear^50^. We speculate that genotoxic crosslinking agents do not induce apoptosis in vivo because, contrary to the situation in culture, most epidermal cells are not cycling, so replication forks will only rarely collide with the crosslinks to activate the damage repair pathway. Other transcription-coupled repair mechanisms, however, might protect the cells and enable them, after sufficient time, to begin proliferation. Although we find that the in vivo responses to DNA damage ultimately resolve over time, plasticity and hyperplasia coupled with DNA damage is an established mechanism for tumor initiation^51^. Platinum compounds remain widely used in cancer chemotherapy for adjuvant or neoadjuvant treatment and metastatic disease, so our results raise significant concerns about unanticipated side-effects.

## ACKNOWLEDGEMENTS

We would like to thank members of the Macara and Lannigan labs for their insightful feedback throughout this project. This work was supported by NCI/NIH grants R35CA132898 to IM, F32CA21374 to LS and T32CA119925 to LS, as well as American Cancer Society grant PF-18-007-01-CCG to LS.

## MATERIALS AND METHODS

### Mouse lines

The mouse lines acquired from Jackson Laboratory (Bar Harbor, ME) for this study include: K14-rtTA (JAX stock # 008099)^16^, (tetO)7-Cre (“TetO-Cre”) (Jax stock # 006234)^17^, Ai9 (“tdTomato”) (JAX stock # 007909)^18^, K8-CreER (JAX stock # 017947)^1^, NSG (JAX stock # 005557)^34, 35^, Rosa26-DTA176 (“DTA”) (JAX stock # 010527)^52^, Col1a2-CreER (JAX stock # 029567)^40^, and R26R-Confetti (“Confetti”) (JAX stock # 013731)^25, 26^. C3H mice (Strain code 025) were acquired from Charles River Laboratories (Wilmington, MA). All mouse experiments were performed with approval from the Vanderbilt Institutional Animal Care and Use Committee. For BrdU treatment, mice were injected with 10mg/kg BrdU (MilliporeSigma, Burlington, MA) and sacrificed after 3 hr.

### Mammary surgery and drug treatment

Female K14-rtTA; TetO-Cre; tdTomato or K8-CreER; tdTomato mice (21-28 days old, at puberty onset) were anesthetized and small circular incisions were made around the 4^th^ pair of nipples to allow access to the underlying mammary fat pads. Fat pads were injected with 4mM cisplatin (left) or PBS vehicle (right). For saturation studies, mice were immediately put on Dox Diet following surgery (Bio-Serv, Flemington, NJ) and maintained for six weeks before sacrifice and sample collection (see Timeline, Fig.1B).

### Epidermal surgery and drug treatment

Following anesthesia administration, a razor was used to shave and make superficial abrasions (referred to throughout the text as “mechanical wounding” or “scrape”) on the backskin of 3-6 wk old adult C3H and K14-rtTA; TetO-Cre; tdTomato/Confetti mice. Cisplatin (4mM) (BioVision, Inc., Milpitas, CA) or PBS (vehicle) was then spread (“painted”) across the shaven region. For intradermal injections, a shaven region of backskin was pinched with forceps and a 26G needle was used to superficially inject 30uL of drug or vehicle into multiple regions within the shaved area.

### Cryosectioning, immunohistochemical staining and image acquisition

Mammary and epidermal tissue samples were cryoembedded in O.C.T. (Fisher Scientific, Hampton, NH). A Leica C1950 cryostat was used to generate 50 *μ*m sections of mammary tissue and 12 *μ*m sections of epidermal tissue. Tissue sections were fixed for 8 min (epidermis) or 15 min (mammary) in 4% paraformaldehyde. BrdU-injected tissue was subjected to an additional 1 hr 2N HCl fixation at 37°C prior to BrdU staining. The primary antibodies used in this study included: chicken α-keratin 14, rabbit α-keratin 10 and rabbit α-keratin 6 (Biolegend, San Diego, CA), rat α-keratin 8 (Developmental Studies Hybridoma Bank, Iowa City, Iowa), rat α-BrdU (Abcam, Cambridge, UK), rat α-E-cadherin (ThermoFisher Scientific, Waltham, MA), rabbit α-p63 (Cell Signaling Technology, Danvers, MA), and mouse α-phospho-histone H2A.X (MilliporeSigma, Burlington, MA). Secondary antibodies included: Alexa Fluor 488, 594 and 647 conjugates of anti-chicken, -mouse, -rat and -rabbit (ThermoFisher Scientific, Waltham, MA). Images were acquired using a Nikon A1R line scanning confocal microscope with a spectral unmixing module (applied for confetti imaging), 20X/0.75 numerical aperture (NA), 40X/1.30 NA and 60X/1.40 NA Plan Apochromat objectives, Type B Immersion oil (Cargille Laboratories, Cedar Grove, NJ) and Nikon NIS-Elements imaging software. Fiji (ImageJ) software was utilized for post-acquisition processing.

### Fibroblast transplantation assay

Adult C3H mouse (3-6 wk donors) backskin was shaven, scraped and painted (as described above) with 4mM cisplatin or PBS vehicle and left for three days. Mice were then sacrificed and backskin was isolated, processed and maintained at 4°C overnight in 0.25% Trypsin-EDTA (ThermoFisher Scientific, Waltham, MA), as previously described^53^. Fibroblasts were isolated the following day, as previously described^54^. Fibroblasts were then injected into the dermis of adult C3H mice (3-6 wk syngeneic recipients). A total of 240,000 fibroblasts were injected per backskin (ten 30 *μ*l intradermal injections of 24,000 cells spaced along different areas of the shaved region). Mice were left for three days before sacrifice and tissue analysis (see Timeline, Fig.4H).

### Fibroblast ablation assay

The backskin of Col1a2-CreER; DTA mice was shaven, scraped and painted with 4-Hydroxytamoxifen (4OHT) (Sigma-Aldrich, St. Louis, MO) or 100% ethanol (EtOH, vehicle) according to the timeline in Fig.4A. Following four consecutive days of 4OHT or EtOH treatment, the same shaven backskin region was re-scraped and repainted with 4OHT or EtOH in addition to 4mM cisplatin. Mice were then left for seven days before sacrifice and tissue analysis.

### Pulse-chase experiments

For epidermal pulse-chase experiments, adult K14-rtTA; TetO-Cre; tdTomato/Confetti mice (3-6 wk old) were injected with a total of 5 mg doxycycline over 5 days (dox pulses: day 1- 2mg, day 3- 2 mg, day 5- 1 mg) while also on Dox Diet (see Timeline, Fig.2C). A region of backskin was shaven, scraped and painted on day 7, and mice were switched onto a regular diet (chase) for 7 days before sacrifice and tissue analysis. For mammary pulse-chase experiments, K14-rtTA; TetO-Cre; tdTomato mice were injected with two pulses of 4.5mg doxycycline, the first at 4 wks old and the second at 7 wks old (Timeline, Fig.S1S).

### Cell culture treatments

EpH4 cells at 50% confluency (the same cell numbers were plated for each condition) were treated with 1 *μ*M or 3 *μ*M Cisplatin or Mitomycin C (or PBS and dH_2_0 vehicles, respectively). After 24 hr incubation at 37°C and 5% CO_2_, drugs were washed off, cells were split, and 1 mL of each was plated into a separate 10 cm dish. Cells were imaged and counted 48 hrs later, and the number of cells on experimental vs vehicle plates was quantified. For cell mixing experiments, stably transduced mApple and eGFP EpH4 cells were sorted using a BD 4 laser FACSAria III cell sorter (BD Biosciences, San Jose, CA), treated for 24 hours with 1 *μ*M Cisplatin or PBS vehicle, and then cultured until Cisplatin-treated cells recovered proliferative activity. Cisplatin- and vehicle-treated cells were then mixed at a 1:1 ratio and grown to confluency (~48 hours) before imaging. GFP:mApple ratios were quantified by thresholding each channel, then overlaying onto nuclei ROIs for counting using Nikon Elements software. P-values were generated by comparing each bar with those of “Veh/Veh”.

### Quantifications

Epidermal thicknesses were quantified using Fiji software by measuring from the base of the epidermis (bottommost K14-positive basal cell layer) to the top of the epidermis (topmost K10-positive suprabasal cell layer) (yellow bracketed region, Fig.2F). Neither hair follicle shafts nor infundibular regions of the upper hair follicle were included in the thickness measurements.

### Statistical analyses

Thickness measurements were analyzed using Student’s *t*-tests to compare between vehicle and Cisplatin measurements of the same day, as well as to analyze how measurements within the same treatment group changed over the experimental time period (p < 0.05).

## SUPPLEMENTARY FIGURES

**Figure S1.**
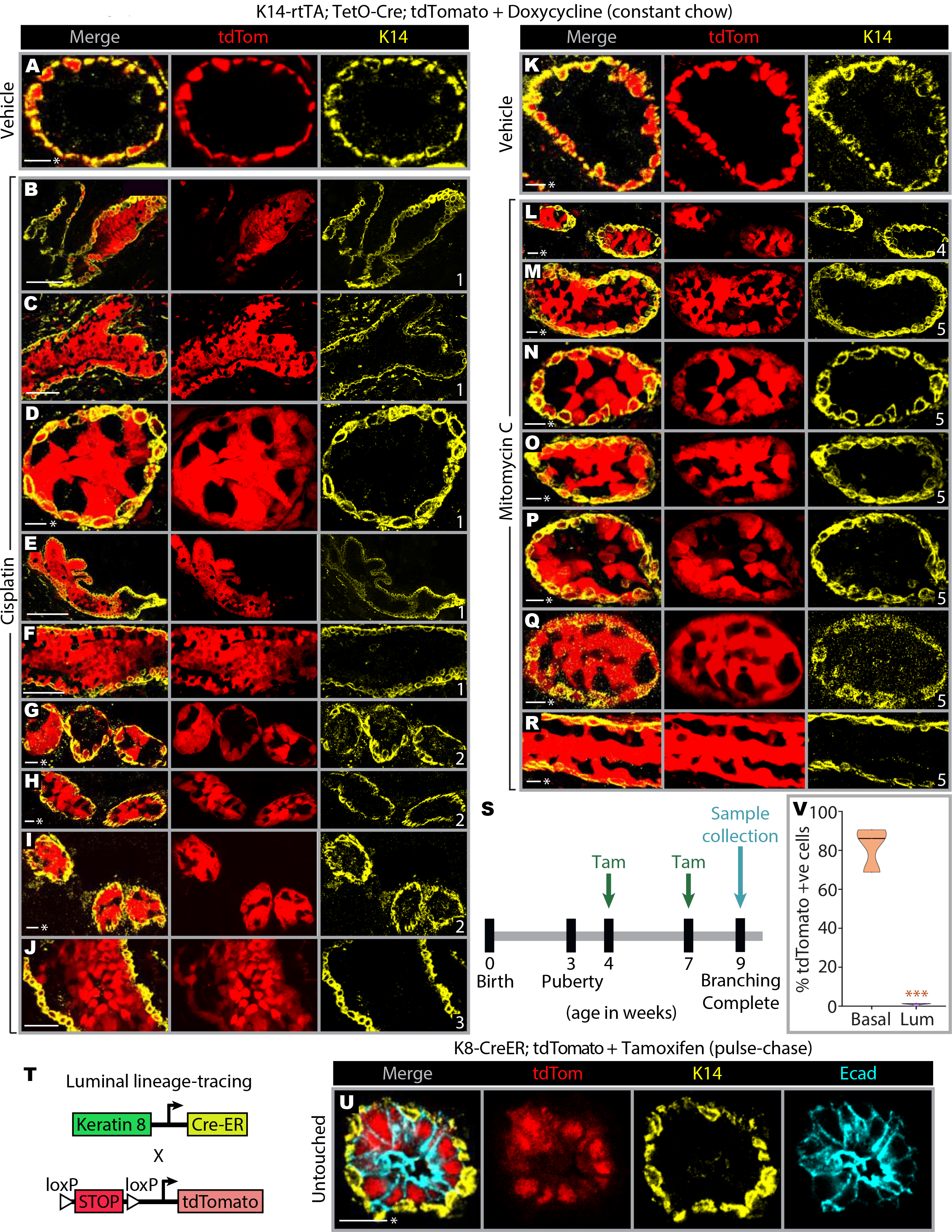
DNA damage enhances cell plasticity and causes luminal filling in mouse mammary glands. A) Representative immunofluorescence image of a mammary cross section from a vehicle (PBS)-treated mouse (Cisplatin control). B-J) Immunofluorescence images of tdTomato-positive luminal cells and luminal filling in the mammary glands of Cisplatin-treated K14-rtTA; TetO-Cre; tdTomato mice, according to the experimental timeline in Fig.1B. Numbers in the bottom right of the K14 channel images indicate the individual mouse from which the image was taken. K) Representative immunofluorescence image of a mammary cross section from a vehicle (dH_2_O)-treated mouse (Mitomycin C control). L-R) Immunofluorescence images of tdTomato-positive luminal cells and luminal filling in the mammary glands of Mitomycin C-treated K14-rtTA; TetO-Cre; tdTomato mice, according to the experimental timeline in Fig.1B (4/4 mice, 100%). S) Experimental timeline of tamoxifen (“tam”)-driven induction of luminal cell labeling. T) Diagram illustrating the two transgenic mouse lines crossed for mammary luminal lineage-tracing. U) Immunofluorescence image showing that unperturbed luminal cells primarily give rise to luminal progeny throughout morphogenesis (2/2 mice, 100%). V) K14-rtTA; TetO-Cre; tdTomato mice were treated with doxycycline according to the timeline in Fig.1B, and tdTomato-positive mammary epithelial cells within the basal and luminal compartments were then quantified by flow cytometry (n=3 mice). ***p < 0.0001. Scale bars are 10 *μ*m (*) or 50 *μ*m.

**Figure S2.**
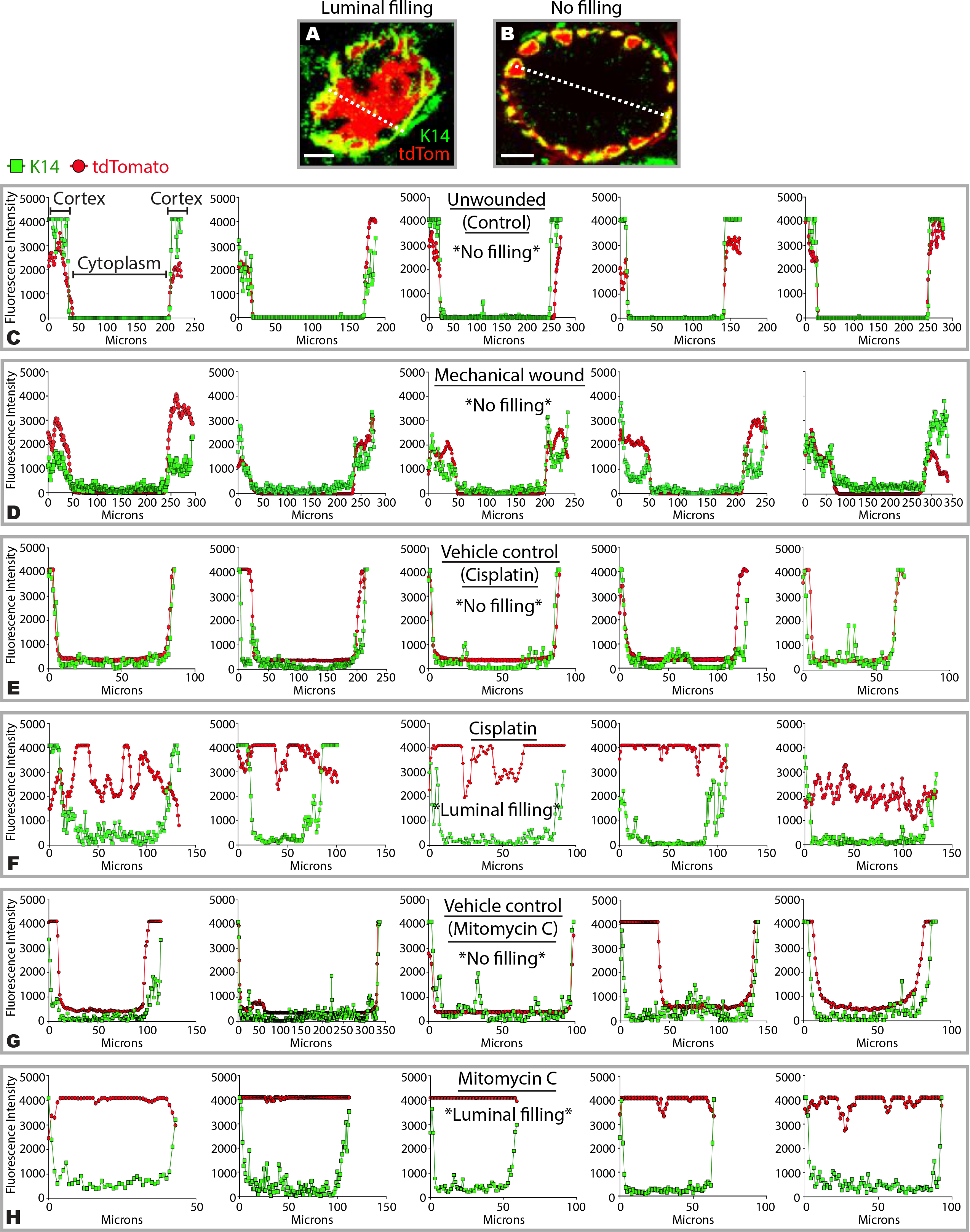
DNA damage promotes luminal filling in mouse mammary glands. A-B) Representative immunofluorescence images of mammary cross sections that exhibit (A) or do not exhibit (B) luminal filling. A line is drawn from the K14+ myoepithelial layer to the opposing myoepithelial layer in order to generate a line scan for assessing the extent of luminal filling across different treatment conditions. C-D) Line scan graphs from five unwounded (C) or wounded (D) mammary cross sections, revealing no luminal filling in either condition. E-F) Line scan graphs from five vehicle (PBS)- (E) or Cisplatin- (F) treated mammary cross sections, revealing no luminal filling and luminal filling, respectively. G-H) Line scan graphs from five vehicle (dH_2_0)- (G) or Mitomycin C- (H) treated mammary cross sections, revealing no luminal filling and luminal filling, respectively. Scale bars are 10 *μ*m.

**Figure S3.**
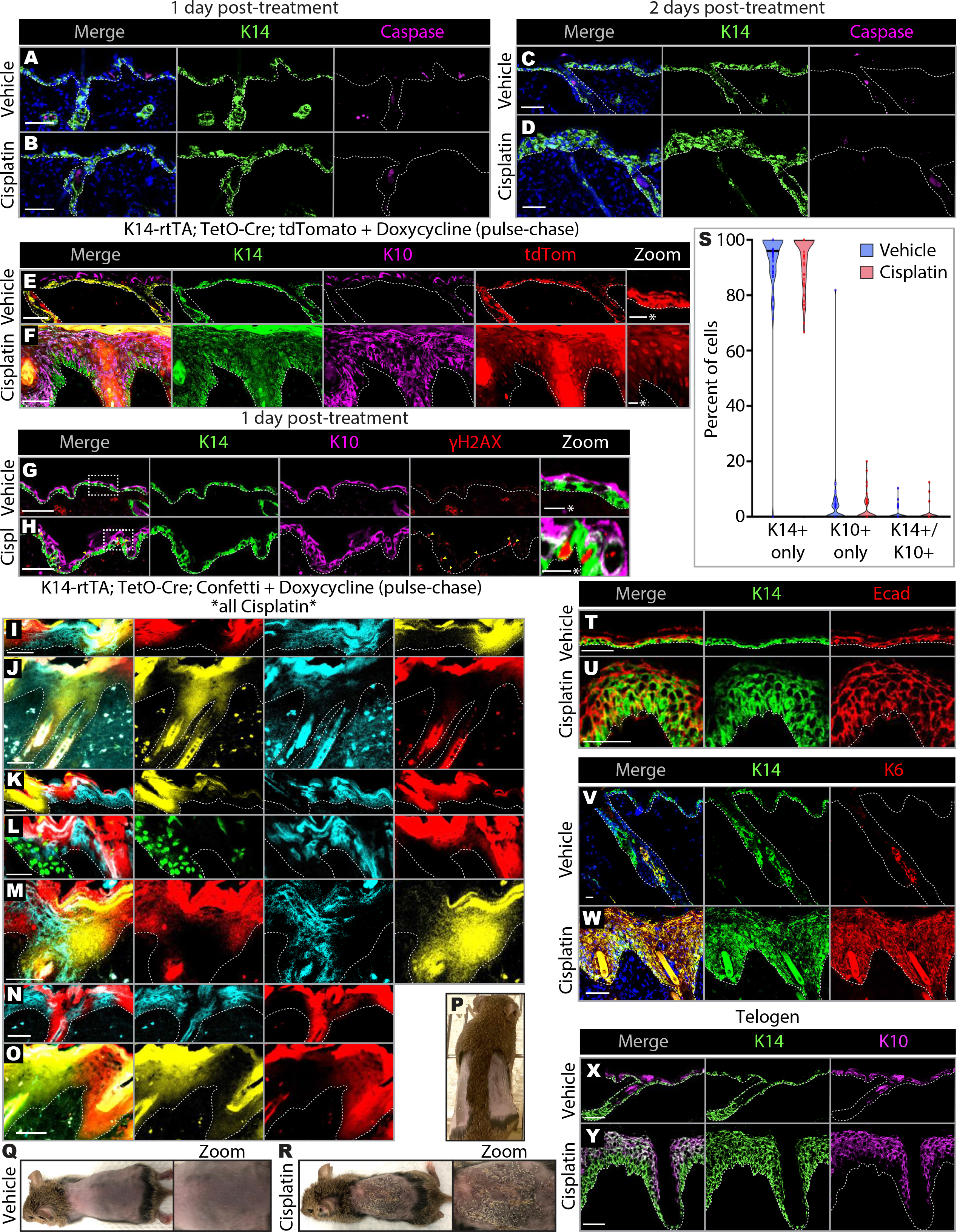
Epidermal tissue maintains epithelial character and does not undergo apoptosis following DNA damage in vivo. A-D) Immunofluorescence images showing cleaved caspase-3 staining of wild-type C3H epidermal tissue, revealing that epithelial and dermal cells do not apoptose in response to mechanical wounding 1 day (A) or 2 days (C) after treatment, nor in response to Cisplatin 1 day (B) or 2 days (D) after treatment. E-F) Immunofluorescence images of epidermis from pulse-chase lineage-tracing in K14-rtTA; TetO-Cre; tdTomato mice, revealing that all suprabasal layers are generated from hyper-proliferative basal cells following Cisplatin treatment, according to the experimental timeline in Fig.2C (n=3 mice). G-H) Immunofluorescence images of γH2AX staining of wild-type C3H epidermal tissue 1 day after Cisplatin treatment reveals positive nuclear signal in Cisplatin-treated basal cells and dermis (H), unlike γH2AX-negative control tissue (G) (note - the red dermal signal in (G) is non-specific and non-nuclear). Yellow arrowheads in (H) indicate γH2AX-positive basal cell nuclei. I-O) Additional immunofluorescence images of Cisplatin-treated K14-rtTA; TetO-Cre; Confetti epidermal tissue, accompanying the timeline and data presented in Fig.2C, J-K. P) Photograph of the two distinct shaven backskin regions (for vehicle and experimental applications) separated by a tuft of hair (represented by a dotted line in Figs.2B, 3C). Q-R) Photographs of gross backskin appearance following vehicle (Q) or Cisplatin (R) treatment. S) Violin plots showing results from treating primary epidermal cells isolated from neonatal mice with vehicle (PBS) or Cisplatin for 24 hours. Cells were assessed for lineage infidelity by quantifying the percentage of cells exhibiting K14/K10 double-positive staining (n=2 experimental replicates). T-U) Immunofluorescence images showing E-cadherin staining of wild-type C3H epidermal tissue, revealing that both vehicle- (T) and Cisplatin- (U) treated cells maintain their epithelial character based on robust cortical E-cadherin localization. V-W) Immunofluorescence images of Keratin 6 staining of wild-type C3H epidermal tissue reveals that, unlike vehicle (V), Cisplatin-treated cells (W) are under stress, based on the presence of positive staining in the interfollicular epidermis that is normally absent from unperturbed tissue. X-Y) Immunofluorescence images of four month old wild-type C3H mouse backskin in the telogen (resting) phase of the hair cycle that was treated with vehicle (X) or Cisplatin (Y) (n=2 mice). Cisplatin-treated telogen backskin responded similarly to Cisplatin-treated backskin in the anagen (growth) phase (all other epidermal data throughout this manuscript were generated from anagen backskin). This suggests that the epidermal Cisplatin response is independent of hair cycle phase. Unless otherwise specified, all image panels in this figure correspond to the experimental timeline presented in Fig.2A. Scale bars are 10 *μ*m (*) or 50 *μ*m.

**Figure S4.**
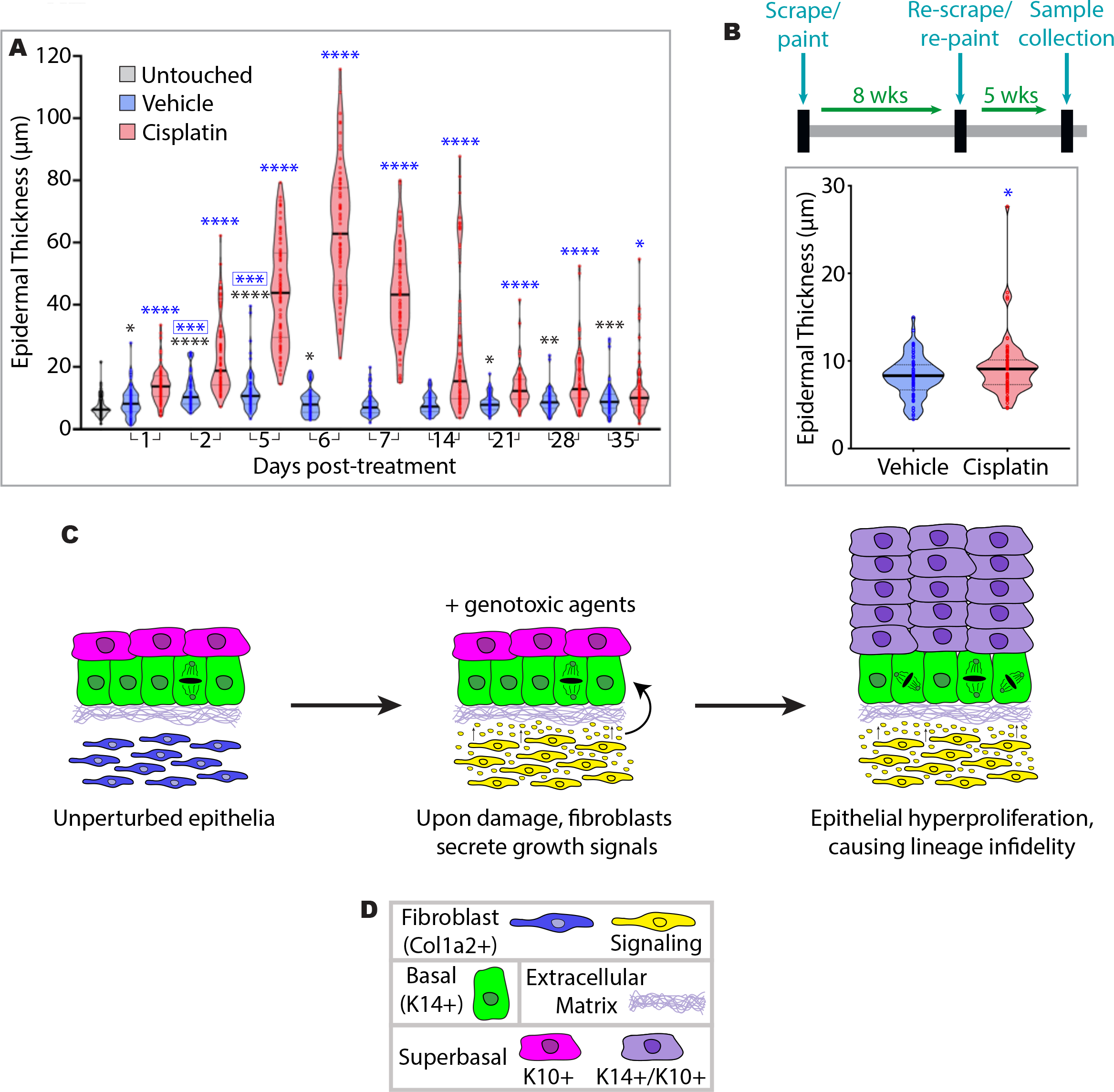
DNA-damaged epidermal tissue resolves over time, and the current working model. A) Violin plots of epidermal thickness over 5 weeks following vehicle and Cisplatin treatment (n=3 mice per violin). Black * is vehicle compared to untreated; blue * is Cisplatin compared to same day vehicle; boxed blue * is vehicle compared to day 1 vehicle. *p < 0.05, **p < 0.001, ***p < 0.0001, ****p<0.00001. Note - days 1-7 are identical data to those presented in Fig.2G. B) Experimental time-course and corresponding violin plots, revealing that repeating Cisplatin treatment does not affect the ability of epidermal tissue thickness to resolve back to near vehicle levels over time (n=3 mice per violin). *p < 0.05. C) Working model of the effect of DNA damage on epithelial behavior. Upon DNA damage, fibroblasts secrete signals to epithelia that promote hyper-proliferation, resulting in lineage infidelity. D) Key of cell types and structures in (C).

